# Empowering Beginners in Bioinformatics with ChatGPT

**DOI:** 10.1101/2023.03.07.531414

**Authors:** Evelyn Shue, Li Liu, Bingxin Li, Zifeng Feng, Xin Li, Gangqing Hu

**Affiliations:** Department of Microbiology, Immunology & Cell Biology, West Virginia University, Morgantown, WV, USA; College of Health Solutions, Arizona State University, Phoenix, AZ, USA; Biodesign Institute, Arizona State University, Tempe, AZ, USA; Finance Department, John Chambers College of Business and Economics, West Virginia University, Morgantown, WV, USA; Department of Economics and Finance, The University of Texas at El Paso, El Paso, TX, USA; Lane Department of Computer Science and Electrical Engineering, West Virginia University, Morgantown, WV, USA

## Abstract

The impressive conversational and programming abilities of ChatGPT make it an attractive tool for facilitating the education of bioinformatics data analysis for beginners. In this study, we proposed an iterative model to fine-tune instructions for guiding a ChatGPT in generating code for bioinformatics data analysis tasks. We demonstrated the feasibility of the model by applying it to various bioinformatics topics. Additionally, we discussed practical considerations and limitations regarding the use of the model in chatbot-aided bioinformatics education.

## Introduction

The recent emergence of large language modes (LLMs), such as ChatGPT, has sparked great interests in their potential use in facilitating programming-heavy data analysis, such as bioinformatics. With its remarkable conversational and programming abilities [1], ChatGPT holds great promise for helping students overcome the programming hurdle. However, as an advanced artificial intelligence (AI) system, the behavior of a chatbot heavily relies on prompts provided by human operators. To fully harness this potential to assist scientific data analysis, prompts used to instruct a chatbot must be carefully crafted so that responses from the chatbot are valid and results are robust.

### The OPTICAL model

Inspired by adaptive learning in educational literature [2], we proposed the OPTICAL model to facilitate chatbot-aided scientific data analysis: Optimization of Prompts Through Iterative Conversation and Assessment with an LLM chatbot (Figure S1). The model involves a series of iterative steps to improve communication with a chatbot for scientific data analysis and enhance students’ learning outcomes. Students will first review the scientific question, analysis task, computational methods, and expected outputs. They will receive guidance on creating a draft prompt describing the data analysis task at various levels of details. Students then converse with the chatbot by inputting the prompt to generate code and execute the chatbot-produced code. In the event error messages are issued after running the code, students must evaluate the error messages and determine the best way to proceed, such as instructing the chatbot to revise the code given the error messages or debugging the code manually. This process iterates until the code no longer issues errors and outputs a result for critical assessment. For an unexpected result, the prompt will be reevaluated and refined, repeating until the expected result is obtained. At the end of the session, students should reflect on the entire communication process and review the code to identify any missing details to be added to the initial prompts. This step may require students to consult relevant manuals and summarize the analytic methods to ensure accuracy and reproducibility. In the end, the iteration and final review expect to yield clear, focused, and concise prompts, as well as a reference code for the desired data analysis.

### Case studies

As a proof-of-concept, we applied the model to three case studies from different topics of bioinformatics and summarized the findings as follows.

#### Short sequencing reads alignment and visual inspection in next generation sequencing analysis

Alignment, the process of determining genomic positions of short sequencing reads, is a fundamental step in deep sequencing data analyses. In this case study, we visually inspected the quality of chromatin immunoprecipitation followed by sequencing dataset generated by Encyclopedia of DNA Elements [3] (see Table S1; Figure S2a). To this end, we instructed ChatGPT to generate code to align the short reads to the human reference genome and summarize the alignments into count numbers across the genome. The results were visually assessed by loading the summarized alignments to Integrative Genomics Viewer [4]. The initial prompts included details of the analyses and bioinformatics tools. The interaction involved two iterations where we instructed the chatbot to handle error messages generated from running the code. Analyzing the final code and reflecting on the entire interactions identified details missing from the initial prompts.

#### Phylogeny inference by DNA sequences in molecular evolution

Phylogenetic inference is an essential yet challenging subject in molecular biology curricula. To demonstrate how ChatGPT can assist students in phylogenetic analyses, we asked the chatbot to generate R code to build a phylogenetic tree for nine species (see Table S2; Figure S2b). This case study started with multiple alignments of protein-coding sequences of the TP53 tumor suppressor gene. The initial prompts included a description of major steps to build an unrooted tree. With two rounds of iterations and human feedback on error messages from running the code, the chatbot wrote workable code to generate a reasonable unrooted phylogenetic tree. We then instructed the chatbot to use a designated species as an outgroup to root the tree. For this complicated task, the chatbot failed to find a valid solution and began to make up functions that did not exist. In this situation, human intervention was required to correct the code after multiple failed iterations.

#### Robust circles fitting in computer vision

Biomedical imaging captures and examines images in biotechnology and medicine. Human vision systems can recognize many different objects in various challenging environments, such as cluttered backgrounds and extreme poses. Circles are arguably the simplest geometric objects for training a computer to recognize. However, instructing a computer to fit circles is a nontrivial task that requires advanced mathematical preparation and proficient computer programming skills. More importantly, students often face the challenge of decomposing a complex problem into several more manageable sub-problems (i.e., a divide-and-conquer approach). In contrast to the previous case studies, this one illustrated a scenario in which describing all analyses in one prompt failed to generate workable code. Instead, we demonstrated how ChatGPT could serve as a virtual teaching assistant to teach the divide-and-conquer approach to a student (see Table S3; Figure S2c). Using a chain-of-thought (CoT) prompt, the student can gradually learn how to solve more and more challenging circle fitting problems, from single to multiple, from clean to noisy observation, and from analytical to numerical, as well as how to incorporate Bayesian priors into the solution algorithms. The results of this experiment was a sophisticated circle-fitting algorithm that cannot be obtained through iterations but could be achieved through CoT prompt design [5].

### Practical considerations

Our firsthand experience with ChatGPT has identified several practical considerations for implementing the OPTICAL model in education settings. To streamline the iteration process, it is crucial to clearly define how the chatbot should respond to prompts, such as acting as an expert in bioinformatics and being proficient in a designated language, outputting code with a minimal number of lines, and resetting the thread upon request. Knowing how to read and interpret code is one focus of the chatbot-assisted training. Other essential skills include using basic commands, reading user manuals of coding packages, and installing software.

Merely using the chatbot as a code-generating tool may limit creative thinking. Therefore, reviewing the code at the end of each session is just as important as optimizing the prompts. At this stage, the focus is on being familiar with the code and identifying missing details in the initial prompts. A well-crafted prompt should be robust across different chat sessions by yielding consistent results. Beginners may start communicating with the chatbot using natural language. When transitioning into intermediate or advanced levels, they may include code in the prompts to keep the chatbot on track.

In addition to overcoming the programming hurdle, another significant impact of the model is to enhance students’ abilities in critical thinking and evaluation of the chatbot’s response. We have observed instances where ChatGPT produced erroneous functions, misused certain options, and faked author name of a package. While many of these errors can be detected by running the code, it is crucial to cross-reference with the manual to ensure a precise understanding of the functions, options, and chatbot’s comments on code, as well as an accurate description of the methods.

A successful application of our model to a specific bioinformatics data analysis task is expected to generate a set of prompts and their associated code, which we refer to as a reference code. While the results from running the reference code should be deterministic if not involving any nondeterministic algorithm, ensuring the robustness of the prompts requires a systematic validation approach. One possible method is to have multiple users run the prompts in new chat sessions and compare the results to the reference one. In research projects where a reference is often absent, the results should be validated through external methods such as literature and existing or additional experiments the same as conventional data analysis. To ensure reproducibility, the prompts, code, and input files used for the project should be made publicly accessible upon publication.

Relevant to the robustness, the same prompts may not generate the same code in a new session. The uncertainty may result from the existence of multiple solutions to the same question or ambiguities/missing details in the prompts, giving the chatbot flexibility to make choices. Educators should be mindful of these uncertainties to control for their disruptions during lectures. On the other side, the uncertainties offer great opportunities for training critical and creative thinking. For example, by comparing new code to the reference, students may learn alternative solutions that improve their bioinformatics skills. Moreover, uncertainties arising from ambiguities in the prompts provide an excellent chance for further refinement through iterations. Nevertheless, novice students must be informed of these uncertainties and potential solutions such as adjusting the temperature setting that controls the chatbot’s response randomness to reduce their anxiety, making chatbot-assisted learning an enjoyable experience.

We vision that a repository of well-defined prompts for typical bioinformatics tasks, along with sample inputs, reference chatbot code, and expected results, would be immensely valuable to beginners. Familiarizing themselves with sample prompts can serve as a steppingstone for students to improve their ability to customize prompts with greater specificity to fit their evolving requirements. The repository also serves as a platform for the community to further validate the robustness of the prompts. However, the challenge remains that prompt engineering tailored for chatbot-aided bioinformatics data analysis in biomedical research, or the broader health science is just an emerging field of research.

### Limitations and Perspectives

The OPTICAL model, like any other model, has its limitations that need to be addressed. The prompt-optimization iteration may not effectively converge to produce a valid solution without in-depth human intervention, especially for advanced data analysis with customized code. Meanwhile, the iterative model may not apply to problems that can be broken down into smaller subproblems that resemble the original problem (known as recursion). An alternative solution is to use the chain-of-thought (CoT) prompt design, as demonstrated in the computer vision case study.

In addition to bioinformatics, case studies in economics and finance (Tables S4 and S5; Figures S2d and S2e) supported a potential extension of the model to scientific data analysis in other disciplines. However, all these case studies were mainly performed by experienced researchers in scientific data analysis with teaching experience. To examine the impact on students, controlled experiments conducted in a classroom setting are needed. Further research is necessary to evaluate the effectiveness of the OPTICAL model in improving students’ learning outcomes in data analysis relative to traditional lecturing. Lastly, an extension of the model to innovative bioinformatics research remains to be explored. As an ongoing effort, we are working on applying the model to new algorithm developments in single-cell gene expression data analysis.

Disadvantaged students beginning to learn bioinformatics face additional barriers like limited access to tutoring services, lack of interactions with instructors, difficulty forming study groups with academically advanced peers, and so on. With a positive outlook, we argue that ChatGPT has the potential to address this knowledge-dissemination disparity. In this human-AI integrative learning process, students may serve as mentors to guide the chatbot in bioinformatics data analysis and, at the same time, learn coding skills from the chatbot. Students may interact with ChatGPT as if studying with a peer that responds instantaneously. This is extremely valuable for academically challenged students who often struggle to find capable peers.

## Conclusions

In conclusion, the OPTICAL model represents a promising step forward in chatbot-aided education in bioinformatics data analysis for beginners. While the concept of ChatGPT-aided education is relatively new [6], our case studies from different disciplines demonstrated ChatGPT’s potential to enhance students’ coding skills and critical creative thinking. Such benefits of practicing bioinformatics with a chatbot are likely to extend from the classroom to a lifelong learning experience, especially for beginners.

## SUPPLEMENTARY MATERIALS

Figure S1: The OPTICAL model for LLM chatbot-assisted scientific data analysis

Figure S2: Summary of case studies applying the OPTICAL model to chatbot-assisted data analysis in five distinctive fields

Table S1: Case study for short sequencing reads alignment and visual inspection

Table S2: Case study for phylogeny inference by DNA sequences

Table S3: Case study for robust circles fitting

Table S4: Case study for household income vs. high school graduation rates

Table S5: Case study for time series analysis of trading data

## ACKNOWLEDGEMENTS

NIH-NIGMS grants P20 GM103434, U54 GM-104942, and 1P20 GM121322 to GH; NIH-NLM grant R01LM013438 to LL. We thank Dr. Jackie J.D. Han from Peking University, Dr. Heather Henderson from West Virginia University, and Dr. Dong Xu from University of Missouri for insightful discussions. The writing was polished by ChatGPT.

## Notes

### Competing Interest Statement

The authors have declared no competing interest.

