## Supplementary figures and images for "Empowering Beginners in Bioinformatics with ChatGPT"

### FigureS1

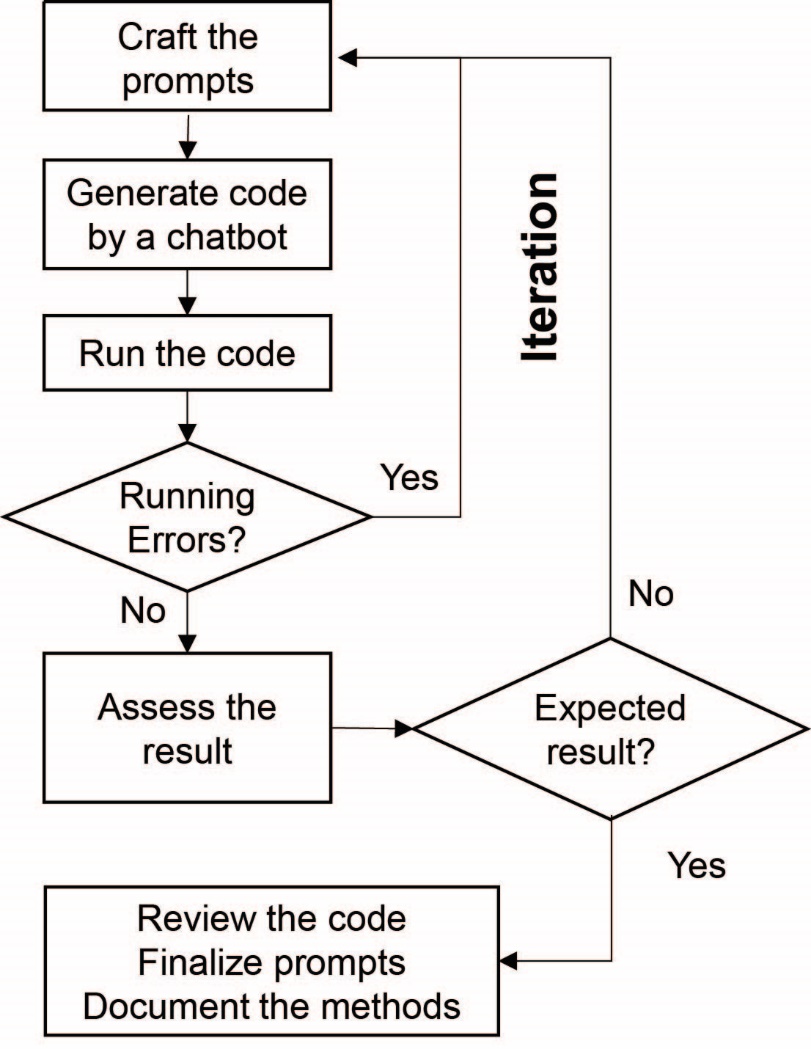

**Figure S1: The OPTICAL model for LLM chatbot-assisted scientific data analysis**
