## Supplementary material for "Empowering Beginners in Bioinformatics with ChatGPT": FigureS2

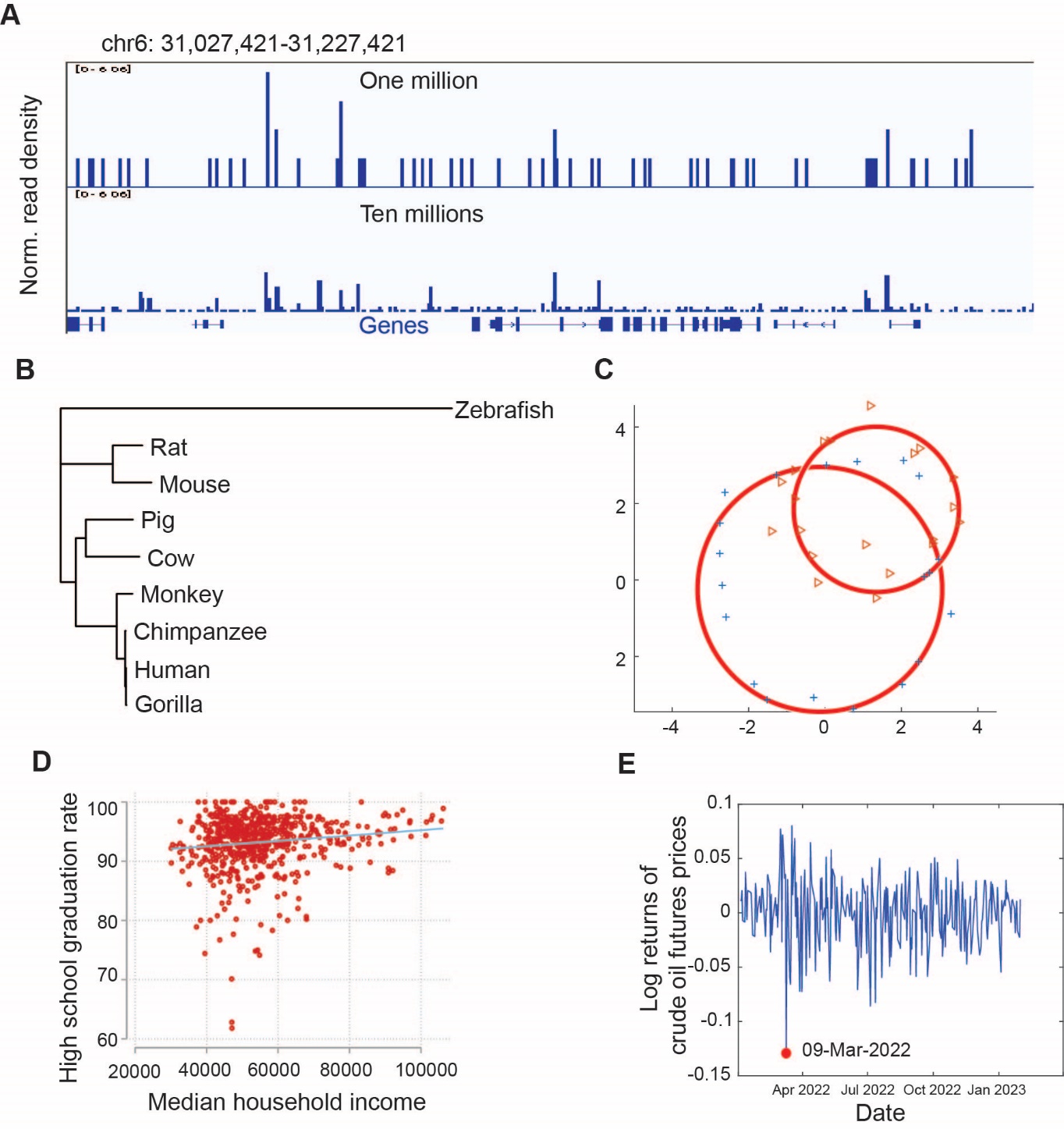

**Figure S2: Summary of case studies applying the OPTICAL model to chatbot-assisted data analysis in five distinctive fields.** (A) Bioinformatics: visualization of ChIP-Seq read density aligned to a 200 KB genomic region in chromosome six at two sequencing depths (one million vs ten million) for transcription factor CTCF in human embryonic stem cell line H1 from ENCODE. (B) Molecular evolution: phylogeny inference of nine species based on protein-coding sequence of TP53. (C) Computer vision: simultaneous fitting of multiple circles to noisy observation. Note that one circle’s inliers behave like another circle’s outliers. (D) Economics: Scatter plot of high school graduation rate vs. median household income with regression line. (E) Finance: Time series plot of log returns of crude oil future prices; Full red circle indicated the lowest returns and the corresponding date.
