## Supplementary material for "Empowering Beginners in Bioinformatics with ChatGPT": TableS1

**OPTICAL CASE STUDY: Short Sequencing Reads Alignment and Visualization**

**Scientific question, analysis methods, expected outputs**

ChIP-seq is a technique used to measure transcription factor (TF) binding by sequencing DNA fragments that have been pulled down by an antibody specific to the TF. The number of reads sequenced at a genomic region is proportional to the likelihood of the TF binding to that region. In this session, we will instruct ChatGPT to align the sequencing reads back to a reference genome and generate a file in bigwig format for visualizing the distribution of the reads across the genome.

We generated two sequencing files of different depths for a CTCF ChIP-seq library in human ESC H1 cell line from ENCODE, one containing one million reads and the other containing ten million reads. We explored the impact of sequencing depth on TF prediction for the CTCF protein. Results will be compared to the visualization of the distribution generated by ENCODE.

Tools to be used: Bowtie2, samtools, and deepTools.

**Coding language(s)**

Bash

**DEFINE CHATBOT’S BEHAVIORS**

Act as an experienced bioinformatician proficient in ChIP-Seq data analysis, you will assist me by writing code with number of lines as minimal as possible. Rest the thread if asked to. Reply “YES” if understand.

**ITERATION 0**

**Prompts**

I have two fastq files in current folder from single-end sequencing of a ChIP-Seq library: ENCFF000AVS_1m.fastq.gz, and ENCFF000AVS_10m.fastq.gz. For each fastq file, align reads to the human reference genome, save to bam file, and then covert it to bigwig file. Tools to use: bowtie2, samtools, and deepTools. The index for bowtie2 is in the folder “../data/indx/bowtie2_whole_genome/” with “hg38” as the prefix. Use 24 CPU for the alignment. Please draft the code in bash.

**Code**

#!/bin/bash

\# Define variables

INDEX="../data/indx/bowtie2_whole_genome/hg38"

CPUS=24

FASTQ1="ENCFF000AVS_1m.fastq.gz"

FASTQ2="ENCFF000AVS_10m.fastq.gz"

\# Align reads to reference genome using bowtie2

bowtie2 -p $CPUS -x $INDEX -U $FASTQ1 | samtools sort -@ $CPUS -o ENCFF000AVS_1m.bam

bowtie2 -p $CPUS -x $INDEX -U $FASTQ2 | samtools sort -@ $CPUS -o ENCFF000AVS_10m.bam

\# Convert BAM files to bigWig format using deepTools

bamCoverage -b ENCFF000AVS_1m.bam -o ENCFF000AVS_1m.bw --binSize 10 --normalizeUsing CPM

bamCoverage -b ENCFF000AVS_10m.bam -o ENCFF000AVS_10m.bw --binSize 10 --normalizeUsing CPM

**Result**

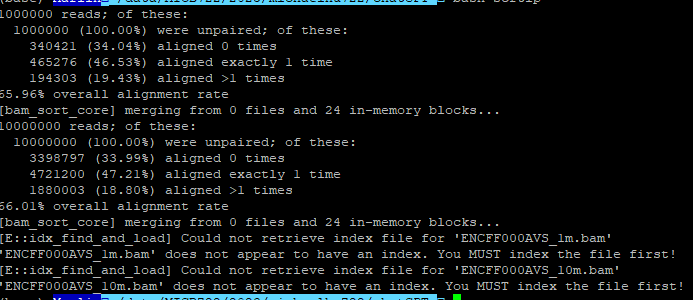

**Assessment***

The alignment worked well. However, error message indicates that bam files are not indexed. This error message was passed to ChatGPT and see its response.

**ITERATION 1**

**Prompts**

[E::idx_find_and_load] Could not retrieve index file for 'ENCFF000AVS_1m.bam'

**Code**

#!/bin/bash

\# Define variables

INDEX="../data/indx/bowtie2_whole_genome/hg38"

CPUS=24

BAM1="ENCFF000AVS_1m.bam"

BAM2="ENCFF000AVS_10m.bam"

\# Sort BAM files using samtools

samtools sort -@ $CPUS -o ENCFF000AVS_1m.sorted.bam $BAM1

samtools sort -@ $CPUS -o ENCFF000AVS_10m.sorted.bam $BAM2

\# Index sorted BAM files using samtools

samtools index ENCFF000AVS_1m.sorted.bam

samtools index ENCFF000AVS_10m.sorted.bam

\# Convert BAM files to bigWig format using deepTools

bamCoverage -b ENCFF000AVS_1m.sorted.bam -o ENCFF000AVS_1m.bw --binSize 10 --normalizeUsing CPM

bamCoverage -b ENCFF000AVS_10m.sorted.bam -o ENCFF000AVS_10m.bw --binSize 10 --normalizeUsing CPM

**Result**

NA

**Assessment***

This time I noticed that “samtools index” was inserted ahead of bamCoverage. Also notice that the code assumed that bowtie alignment was done successfully, and therefore can not be directly applied to address our needs without adding back the code for the alignment.

**ITERATION 2**

**Prompts**

Wait, I saw that you have "*samtools index*" before "*bamcoverage*". Does *bamcoverage* as bam to be sorted before using as input?

**Code**

#!/bin/bash

\# Define variables

INDEX="../data/indx/bowtie2_whole_genome/hg38"

CPUS=24

FASTQ1="ENCFF000AVS_1m.fastq.gz"

FASTQ2="ENCFF000AVS_10m.fastq.gz"

\# Align reads to reference genome using bowtie2

bowtie2 -p $CPUS -x $INDEX -U $FASTQ1 | samtools sort -@ $CPUS -o ENCFF000AVS_1m.sorted.bam

bowtie2 -p $CPUS -x $INDEX -U $FASTQ2 | samtools sort -@ $CPUS -o ENCFF000AVS_10m.sorted.bam

\# Index sorted BAM files using samtools

samtools index ENCFF000AVS_1m.sorted.bam

samtools index ENCFF000AVS_10m.sorted.bam

\# Convert sorted BAM files to bigWig format using deepTools

bamCoverage -b ENCFF000AVS_1m.sorted.bam -o ENCFF000AVS_1m.bw --binSize 10 --normalizeUsing CPM

bamCoverage -b ENCFF000AVS_10m.sorted.bam -o ENCFF000AVS_10m.bw --binSize 10 --normalizeUsing CPM

**Result**

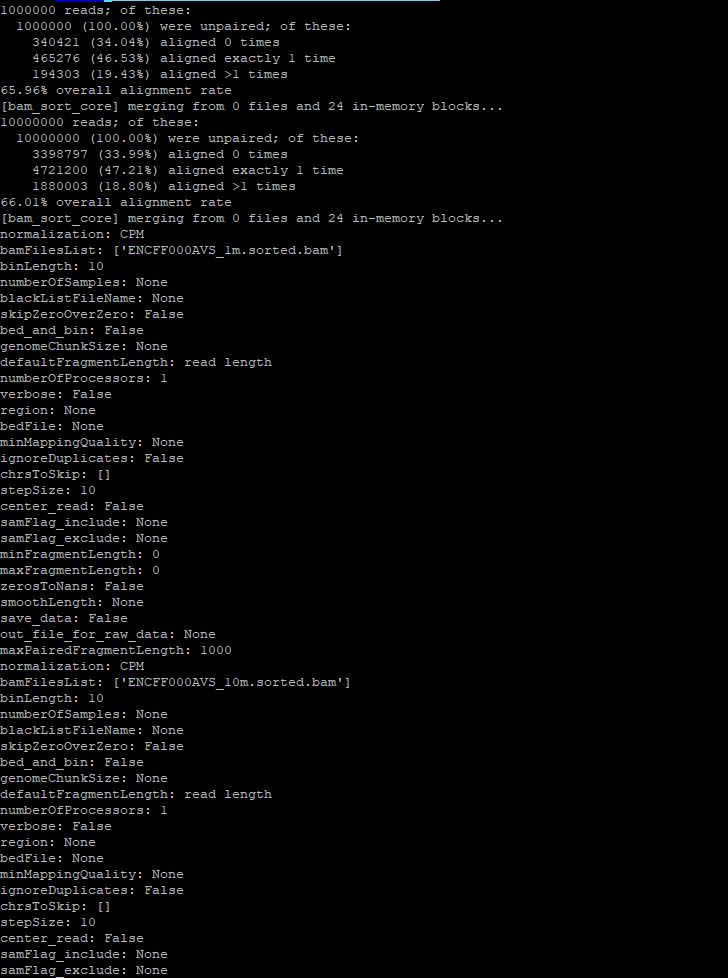

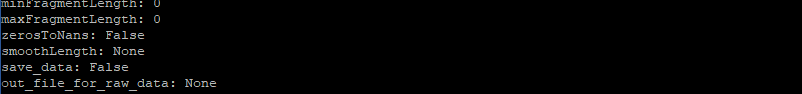

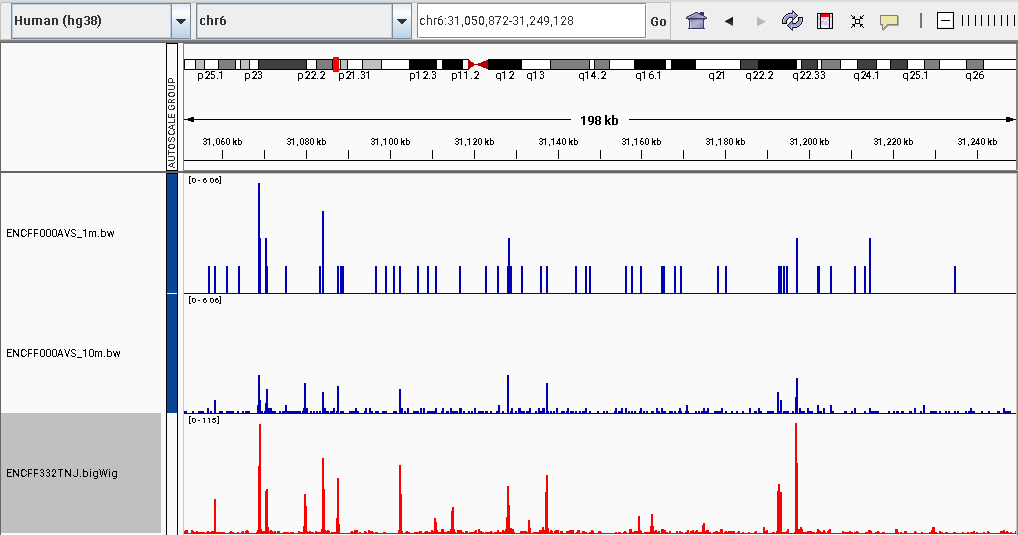

**Figure legend:** Distribution of CTCF ChIP-Seq reads on human H1-hESC across a genomic region of 198 kb. Y-axis for the first two tracks (ENCFF000AVS_1m.bw and ENCFF000AVS_10m.bw) are normalized reads density (CPM: count per million), generated by running codes drafted by chatGPT. Y-axis for the last track represent fold change over control from ENCODE using all the sequencing reads (ENCFF332TNJ.bigWig; 24.6 million).

**Assessment***

The chatbot said that “*bamCoverage* requires that the input BAM file is sorted by genomic position” and generate code to satisfy this requirement. I confirmed that the missing of index error message was issued by *bamCoverage*. The code ran successfully and generated bigwig files for both fastq files.

The bigwig files were uploaded to IGV for visualization. We assessed the results by comparing to the bigwig from ENCODE. The results are generally consistent. The more sequencing depth, the more peaks will be observed.

**WRAPPING UP**

**Review the final codes**

Review the code comments added by ChatGPT and identify key steps, as represented by functions and parameters, that are missing in the initial prompts.

Prompts

I need to insert line-by-line comments to the below code which works well to address the needs for the data analysis task. Wait for my code.

Code with explanations

#!/bin/bash

\# Set the path to the bowtie2 index

INDEX="../data/indx/bowtie2_whole_genome/hg38"

\# Set the number of CPUs to use for alignment

CPUS=24

\# Set the path to the first fastq file

FASTQ1="ENCFF000AVS_1m.fastq.gz"

\# Set the path to the second fastq file

FASTQ2="ENCFF000AVS_10m.fastq.gz"

\# Align the reads from the first fastq file using bowtie2, then sort the resulting SAM file and convert it to BAM using samtools

bowtie2 -p $CPUS -x $INDEX -U $FASTQ1 | samtools sort -@ $CPUS -o ENCFF000AVS_1m.sorted.bam

\# Align the reads from the second fastq file using bowtie2, then sort the resulting SAM file and convert it to BAM using samtools

bowtie2 -p $CPUS -x $INDEX -U $FASTQ2 | samtools sort -@ $CPUS -o ENCFF000AVS_10m.sorted.bam

\# Index the BAM file from the first fastq file using samtools

samtools index ENCFF000AVS_1m.sorted.bam

\# Index the BAM file from the second fastq file using samtools

samtools index ENCFF000AVS_10m.sorted.bam

\# Convert the sorted BAM file from the first fastq file to bigwig using deepTools, with a bin size of 10 and normalized using CPM

bamCoverage -b ENCFF000AVS_1m.sorted.bam -o ENCFF000AVS_1m.bw --binSize 10 --normalizeUsing CPM

\# Convert the sorted BAM file from the second fastq file to bigwig using deepTools, with a bin size of 10 and normalized using CPM

bamCoverage -b ENCFF000AVS_10m.sorted.bam -o ENCFF000AVS_10m.bw --binSize 10 --normalizeUsing CPM

**Missing details in the initial prompts**

1. Index the bam file
2. The bamCoverage has an option “--normalizeUsing" to specify the type of normalization to facilitate comparison between different libraries.

**Crafting the final prompts**

The initial prompts were revised to include the missing details (in bold) discovered during code review.

Act as an experienced bioinformatician proficient in ChIP-Seq data analysis, you will assist me by writing code with number of lines as minimal as possible. Rest the thread if asked to. Reply “YES” if understand.

I have two fastq files in current folder from single-end sequencing of a ChIP-Seq library: ENCFF000AVS_1m.fastq.gz, and ENCFF000AVS_10m.fastq.gz. For each fastq file, align reads to the human reference genome, save to bam file, **index it**, and then covert it to bigwig file with **CPM normalization**. Tools to use: bowtie2, samtools, and deepTools. The index for bowtie2 is in the folder “../data/indx/bowtie2_whole_genome/” with “hg38” as the prefix. Use 24 CPU for the alignment. Please draft the code in bash.

**Summary***

**Aims**

The goal of this session is to optimize the prompts which will guide ChatGPT to generate code for aligning sequencing files (fastq) and then generate a summarized file (bigwig) for the distribution of reads across the genome.

**Materials and Methods**

The original fastq sequencing file was obtained from ENCODE (ENCFF000AVS). In this analysis session, we extracted the first one million reads and the first ten million reads from the original file. In this way, we could tell how sequence depth impacts the finding TF bindings sites even from a visual inspection.

We aligned the sequencing file to the human reference (hg38) genome using Bowtie2 (v 2.4.2)^1^ and converted the output to sorted and indexed BAM files using Samtools (v 1.10)^2^. We then used the *bamCoverage* (v 3.5.1) function from deepTools ^3^ to convert the indexed BAM file to a bigwig file. The bigwig files were visualized by IGV (v2.9.2)^4^.

**Results and Discussions**

Visual inspection of the distribution of CTCF ChIP-Seq reads across the genome revealed a significantly higher number of peaks in the ten million reads dataset compared to the one million reads dataset. Data analysis, such as examining the signal at known CTCF target sites in ESC, conducting motif enrichment analysis, and comparing to other public CTCF ChIP-Seq datasets can be used as additional quality control for the current library.

**Additional Comments***

Students are encouraged to test the robust of these prompts with new ChatGPT sessions. Alternatively, students could drop out certain details in the prompts to check what alternative solutions ChatGPT may come up with.

*The writing has been polished by ChatGPT after an initial human draft
