## Supplementary material for "Empowering Beginners in Bioinformatics with ChatGPT": TableS2

**OPTICAL CASE STUDY: Phylogeny inference based on TP53**

**Description:**

Test how ChatGPT can help students build a phylogenetic tree starting with aligned TP53 gene coding sequences from 9 species.

**Coding language(s):**

R

Prompts to d*efine chatbot’s behaviors*

Act as an experienced bioinformatician proficient in R, you will write code with number of lines as minimal as possible. Rest the thread if asked to. Reply “YES” if understand.

**ITERATION 0**

**Prompts**

You have a multiple alignment file named as tp53.clustal in ClustalW format. Please write R code that can load the file, calculate evolutionary distance, build a NJ tree, and visualize the phylogeny.

**Code**

\# Load the required packages

library(ape)

\# Read in the alignment file

aln <- read.alignment("tp53.clustal", format="clustal")

\# Calculate the evolutionary distance

dist <- dist.dna(aln)

\# Build the NJ tree

tree <- nj(dist)

\# Plot the phylogeny

plot(tree)

**Result**

Failed with an error message.

**ITERATION 1**

**Prompts**

I got an error message complaining "could not find function "read.alignment". Please fix it.

**Code**

\# Load the required packages

library(seqinr)

library(ape)

\# Read in the alignment file

aln <- read.alignment("tp53.clustal", format="clustal")

\# Calculate the evolutionary distance using the Jukes-Cantor distance

dist <- dist.dna(aln, model="JC")

\# Build the NJ tree

tree <- nj(dist)

\# Plot the phylogeny

plot(tree)

**Result**

Failed with an error message.

**ITERATION 2**

**Prompts**

I got a warning message " In dist.dna(aln) : NAs introduced by coercion". Please fix it.

**Code**

\# Load the required packages

library(seqinr)

library(ape)

\# Read in the alignment file

aln <- read.alignment("tp53.clustal", format="clustal")

\# Calculate the evolutionary distance using the Jukes-Cantor distance

dist <- dist.dna(aln, model="JC")

\# Build the NJ tree

tree <- nj(dist)

\# Plot the phylogeny

plot(tree)

**Result**

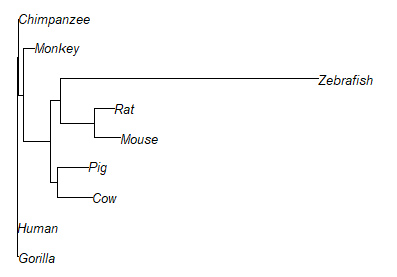

The code displays an unroot tree. Manual editing of the code and instruct ChatGPT to generate a rooted tree.

**ITERATION 3**

**Prompts**

I wrote an R program to read a multiple alignment file named as tp53.clustal in ClustalW format, calculate evolutionary distance, build a NJ tree, and visualize the phylogeny. But I want to root the tree with the Zebrafish sequence as the outgroup. Can you help me revise the R code? Below is my R code.

\# Load the required packages

library(seqinr)

library(ape)

\# Read in the alignment file

aln <- read.alignment("tp53.clustal", format="clustal")

\# Calculate the evolutionary distance

dist <- dist.dna(as.DNAbin(aln))

\# Build the NJ tree

tree <- nj(dist)

\# Plot the phylogeny

plot(tree)

**Codes**

\# Load the required packages

library(seqinr)

library(ape)

\# Read in the alignment file

aln <- read.alignment("tp53.clustal", format="clustal")

\# Identify the index of the Zebrafish sequence in the alignment

zebrafish_idx <- which(aln$species == "Danio_rerio")

\# Calculate the evolutionary distance

dist <- dist.dna(as.DNAbin(aln))

\# Build the NJ tree

tree <- nj(dist, outgroup = zebrafish_idx)

\# Plot the phylogeny

plot(tree)

**Results**

Failed with an error message.

**ITERATION 4**

**Prompts**

I got an error message complaining " Error in nj(dist, outgroup = zebrafish_idx) unused argument (outgroup = zebrafish_idx)". Please fix it.

**Codes**

\# Load the required packages

library(seqinr)

library(ape)

\# Read in the alignment file

aln <- read.alignment("tp53.clustal", format="clustal")

\# Identify the index of the Zebrafish sequence in the alignment

zebrafish_idx <- which(aln$species == "Danio_rerio")

\# Calculate the evolutionary distance

dist <- dist.dna(as.DNAbin(aln))

\# Build the NJ tree

tree <- nj(dist)

\# Root the NJ tree with the Zebrafish sequence as the outgroup

tree_rooted <- root(tree, outgroup = zebrafish_idx)

\# Plot the phylogeny

plot(tree_rooted)

**Results**

Failed with an error message.

**ITERATION 5**

**Prompts**

I got an error message complaining "Error in if (newroot == ROOT) { : argument is of length zero". Please fix it.

**Codes**

\# Load the required packages

library(seqinr)

library(ape)

\# Read in the alignment file

aln <- read.alignment("tp53.clustal", format="clustal")

\# Identify the index of the Zebrafish sequence in the alignment

zebrafish_idx <- which(aln$species == "Danio_rerio")

\# Calculate the evolutionary distance

dist <- dist.dna(as.DNAbin(aln))

\# Build the NJ tree

tree <- nj(dist)

\# Root the NJ tree with the Zebrafish sequence as the outgroup

tree_rooted <- root(tree, outgroup = zebrafish_idx)

\# Plot the phylogeny

plot(tree_rooted)

**Results**

Failed with an error message.

**Manual editing of the code.**

**The code displays a correctly rooted tree.**

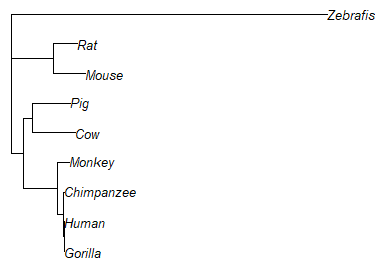

**WRAPPING UP**

**Final codes with explanation to review:**

Prompt: I created the following R code. Please add inline comments.

library(seqinr)

library(ape)

aln <- read.alignment("tp53.clustal", format = "clustal")

dist_matrix <- dist.dna(as.DNAbin(aln))

zebrafish_index <- which(attr(dist_matrix, 'Labels') == "Zebrafish")

tree <- nj(dist_matrix)

tree_rooted <- root(tree, zebrafish_index)

plot(tree_rooted, cex = 0.8)

\# Load the required packages

library(seqinr)

library(ape)

\# Read in the alignment file

aln <- read.alignment("tp53.clustal", format = "clustal")

\# Calculate the evolutionary distance matrix

dist_matrix <- dist.dna(as.DNAbin(aln))

\# Identify the index of the Zebrafish sequence in the distance matrix labels

zebrafish_index <- which(attr(dist_matrix, 'Labels') == "Zebrafish")

\# Build the NJ tree

tree <- nj(dist_matrix)

\# Root the NJ tree with the Zebrafish sequence

tree_rooted <- root(tree, zebrafish_index)

\# Plot the phylogeny with modified label size

plot(tree_rooted, cex = 0.8)

**Summarize the question, methods, and results**

In this session, we aim to assess how ChatGPT can help beginners to build a phylogenetic tree starting with a set of aligned sequences of the TP53 gene from 9 species. We started with a prompt that specified the analysis steps and asked ChatGPT to generate R code. We then copied the code and tested it in R. If the execution generated an error or a warning, we presented the message to ChatGPT and requested it to fix the problem. If ChatGPT couldn’t fix the problem after two iterations, we manually edited the code.

**Additional Comments**

1. ChatGPT was able to solute easy problems such as calling a function in a library but not loading the library beforehand.
2. ChatGPT struggled to resolve more complicated problems, such as identifying and converting to correct data types, and setting advanced parameters in function calls.
3. ChatGPT sometimes made up functions that did not exist.
