## Supplementary material for "Empowering Beginners in Bioinformatics with ChatGPT": TableS3

**OPTICAL CASE STUDY: Robust Circle Fitting Under Matlab: Automatic Algorithm Development**

**Description**

Circle fitting to computer vision is the fruit-fly problem to genetics. It takes a set of 2D coordinates as the input and tries to fit one or multiple circles to the given data. In model-based approaches, circle fitting is often cast as a nonlinear optimization problem. Analytical solutions only exist in simplified scenario (i.e., single circle and clean data). In more challengeing situations (e.g., multiple circles with outliers), RANSAC-based algorithm has been extensively studied in the literature but its Matlab implementation is often beyond the reach of Matlab beginners.

This case study demonstrates how to use ChatGPT as a virtual tutor to solve the problems of circle fitting with gradually increasing difficulty. By decomposing the challenging research problem into a series of chain-of-thought (CoT) prompts, we can assemble a sophisticated circle fitting algorithm from building blocks (like lego-building).

The four prompts in below will be executed in sequential order

1. I need a matlab function for circle fitting.
2. Can you make this fitting function robust to outliers?
3. Can you extend it to robust fitting of multiple circles?
4. If we know that the radius of all circles are within a range of [a,b], can you incorporate this prior information into the fitting algorithm?

ChatGPT failed to generate workable code when the first three prompts were combined into one, for example: “I need a matlab function for fitting multiple circles, robust to outlier points”. (see section “Additional comments”).

**Coding language(s)**

Matlab

Prompts to d*efine chatbot’s behaviors*

Act as an experienced computer vision researcher working with Matlab, you will write code with a number of lines as minimal as possible.

Reply “Yes” if understood.

### STEP 1

**Prompts**

I need a matlab function for circle fitting.

**Code**

**function [center, radius, residual] = fit_circle(x, y)**

**% Fit a circle to a set of 2D points (x, y).**

**% Return the center and radius of the fitted circle.**

**% Also return the residuals (sum of squared distances between the points**

**% and the fitted circle).**

**%**

**% Inputs:**

**% x, y - the x and y coordinates of the points.**

**%**

**% Outputs:**

**% center - a 2-element vector containing the x and y coordinates of the center of the fitted circle.**

**% radius - the radius of the fitted circle.**

**% residual - the sum of squared distances between the points and the fitted circle.**

**%**

**% Example:**

**% x = [1, 2, 3];**

**% y = [2, 3, 4];**

**% [center, radius, residual] = fit_circle(x, y);**

**%**

**% Author: Zhen Li, adapted from the code by Damien Garcia.**

**x = x(:);**

**y = y(:);**

**N = length(x);**

**X = [x, y, ones(N, 1)];**

**Y = -(x.^2 + y.^2);**

**a = X\Y;**

**center = [-a(1)/2, -a(2)/2];**

**radius = sqrt((a(1)^2 + a(2)^2)/4 - a(3));**

**residual = sum((X*a + Y).^2);**

**end**

**Result**

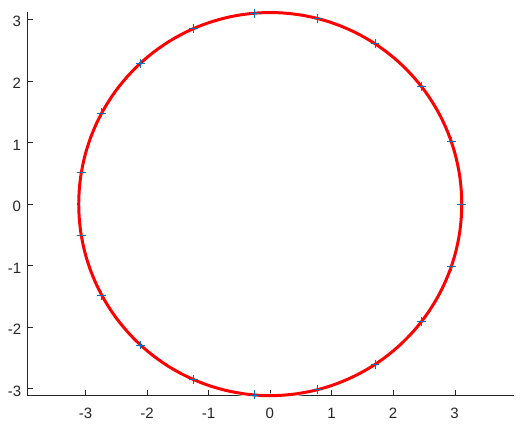

**Assessment**

This is a relatively easy problem to solve.

**STEP 2**

**Prompts**

Can you make this fitting function robust to outliers?

**Code**

function [center, radius, inliers] = ransac_fit_circle(x, y, thresh)

% Fit a circle to a set of 2D points (x, y) using the RANSAC algorithm.

% Return the center and radius of the fitted circle, as well as the indices of the inliers.

%

% Inputs:

% x, y - the x and y coordinates of the points.

% thresh - the maximum allowed distance between a point and the fitted circle.

%

% Outputs:

% center - a 2-element vector containing the x and y coordinates of the center of the fitted circle.

% radius - the radius of the fitted circle.

% inliers - a vector of indices of the inliers (points within the threshold distance).

%

% Example:

% x = [1, 2, 3];

% y = [2, 3, 4];

% thresh = 0.1;

% [center, radius, inliers] = ransac_fit_circle(x, y, thresh);

%

% Author: Zhen Li, adapted from the code by Damien Garcia.

n = length(x);

best_center = [];

best_radius = [];

best_inliers = [];

max_inliers = 0;

for i = 1:1000

% Randomly select 3 points

idx = randsample(n, 3);

X = [x(idx), y(idx)];

[center_i, radius_i] = fit_circle(X(:, 1), X(:, 2));

% Compute the distance between each point and the fitted circle

dist = sqrt((x - center_i(1)).^2 + (y - center_i(2)).^2) - radius_i;

% Identify the inliers (points within the threshold)

inliers_i = abs(dist) < thresh;

% Update the best fitting circle if a larger set of inliers is found

if sum(inliers_i) > max_inliers

best_center = center_i;

best_radius = radius_i;

best_inliers = inliers_i;

max_inliers = sum(inliers_i);

end

end

center = best_center;

radius = best_radius;

inliers = find(best_inliers);

end

**Result**

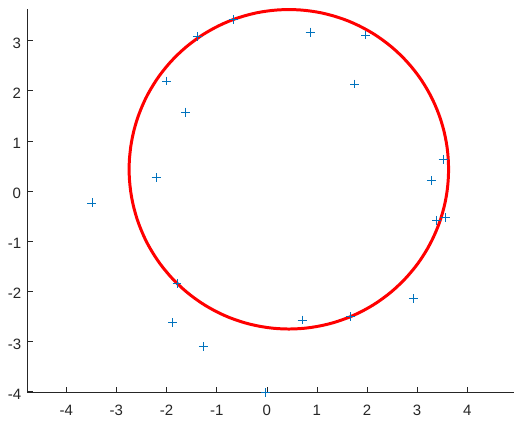

**Assessment**

This is a textbook-based extension of the baseline algorithm.

### STEP 3

**Prompts**

Can you extend it to robust fitting of multiple circles?

**Code**

function [centers, radii, inliers] = ransac_fit_multiple_circles(x, y, thresh, min_points, num_circles)

% Fit multiple circles to a set of 2D points (x, y) using the RANSAC algorithm.

% Return the centers, radii, and indices of the inliers of the fitted circles.

%

% Inputs:

% x, y - the x and y coordinates of the points.

% thresh - the maximum allowed distance between a point and the fitted circles.

% min_points - the minimum number of points required to fit a circle.

% num_circles - the number of circles to fit.

%

% Outputs:

% centers - a num_circles x 2 matrix containing the x and y coordinates of the centers of the fitted circles.

% radii - a num_circles x 1 vector containing the radii of the fitted circles.

% inliers - a cell array of length num_circles, where each element is a vector of indices of the inliers for a fitted circle.

%

% Example:

%	x = [1, 2, 3, 4, 5];

%	y = [2, 3, 4, 5, 6];

%	thresh = 0.1;

%	min_points = 3;

%	num_circles = 2;

%	[centers, radii, inliers] = ransac_fit_multiple_circles(x, y, thresh, min_points, num_circles);

%

% Author: Zhen Li

n = length(x);

points = [x, y];

inliers = cell(num_circles, 1);

centers = zeros(num_circles, 2);

radii = zeros(num_circles, 1);

for i = 1:num_circles

% Fit a circle using RANSAC

[center, radius, inlier_idx] = ransac_fit_circle(x, y, thresh);

centers(i, :) = center;

radii(i) = radius;

inliers{i} = inlier_idx;

% Remove the inliers from the set of points

x(inlier_idx) = [];

y(inlier_idx) = [];

if length(x) < min_points

break;

end

end

end

**Result**

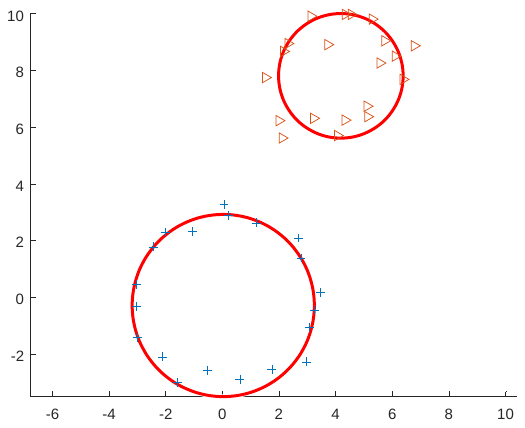

**Assessment**

The code run successfully. The circle fitting results look satisfactory.

### STEP 4

**Prompts**

If we know that the radius of all circles are within a range of [a,b], can you incorporate this prior information into the fitting algorithm?

**Code**

function [centers, radii] = fit_circles_ransac_range(x, y, num_circles, num_iterations, a, b)

% Fit multiple circles to a set of points (x, y) using RANSAC.

% Returns the centers and radii of the fitted circles.

% Set default values for parameters

if nargin < 4

num_iterations = 1000;

end

if nargin < 5

a = 0;

end

if nargin < 6

b = Inf;

end

% Initialize best models

best_centers = zeros(num_circles, 2);

best_radii = zeros(num_circles, 1);

best_num_inliers = zeros(num_circles, 1);

% Loop over circles

for k = 1:num_circles

% Initialize inliers

inlier_indices = 1:length(x);

inlier_x = x;

inlier_y = y;

% Compute distance threshold

if isfinite(b)

r = (a + b) / 2;

R = (b - a) / 2;

threshold = 4 * R^2 / r^2;

else

threshold = 0.01;

end

% Loop over iterations

for i = 1:num_iterations

% Randomly select three points

sample_indices = randperm(length(inlier_x), 3);

sample_x = inlier_x(sample_indices);

sample_y = inlier_y(sample_indices);

% Fit circle to sample points

[sample_center, sample_radius] = fit_circle(sample_x, sample_y);

% Compute distance to circle for all points

distances = abs(sqrt((inlier_x - sample_center(1)).^2 + (inlier_y - sample_center(2)).^2) - sample_radius);

% Count inliers

inlier_indices = find(distances < threshold * sample_radius);

inlier_x = x(inlier_indices);

inlier_y = y(inlier_indices);

num_inliers = length(inlier_indices);

% Check if this is the best model so far

if num_inliers > best_num_inliers(k)

best_centers(k,:) = sample_center;

best_radii(k) = sample_radius;

best_num_inliers(k) = num_inliers;

end

end

% Refit model to all inliers

[centers(k,:), radii(k)] = fit_circle(inlier_x, inlier_y);

% Remove inliers from list of points

x(inlier_indices) = [];

y(inlier_indices) = [];

end

end

**Result**

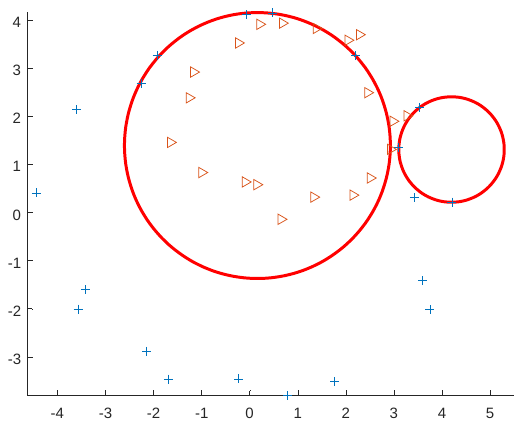

**Assessment**

The code run successfully but the fitting results are not satisfactory.

This is because RANSAC can be trapped by a local minimum during the iterations.

This failure case is an open problem in the literature of CV.

### WRAPPING UP

**Aims**

The goal of this session is to learn how to use Matlab to generate more and more sophisticated circle fitting algorithms.

**Methods**

By providing proper prompts to ChatGPT, the case study demonstrates how beginners could obtain codes from ChatGPT to experiment the circle fitting problem with increasing difficulty levels.

**Results and Discussions**

The Matlab codes generated by ChatGPT work fairly well for simple scenarios such as single circle and multiple circles without noise contamination.

The generated codes start to fail for more challenging situations when multiple circles intersect with each other and have interference from outliers (step 4). This is a well-known weakness of RANSAC-based solution- i.e., the result varies from experiment to experiment due to the randomness of sampling strategy,

The codes obtained from ChatGPT are satisfactory for novice CV researchers. They can use the codes generated by ChatGPT to jump-start their research.

**Additional Comments**

When the first three requirements were combined into one, “I need a matlab function for fitting multiple circles, robust to outlier points”, ChatGPT simply got frozen after outputting the main function. By hitting “regenerate response” button, among the three times of trying, each time ChatGPT tried a different approach but got stuck and could not finish the generation.

This is because for a sophisticated programming task that require calling subroutines (it is called “helper function” under Matlab), ChatGPT did not yet master the art of one function calling another one, including itself (known as recursion).
