## Supplementary material for "Empowering Beginners in Bioinformatics with ChatGPT": TableS4

**OPTICAL CASE STUDY: On Relationship Between Household Income and High School Graduation Rates**

**Description**

This case study aims to explore the relationship between median household income and high school graduation rates in Texas counties from 2020 to 2022, using rigorous regressions and intuitive graphs.

Regression analysis is a long-established and widely used quantitative data analysis technique in the social sciences. It is a compelling statistical procedure that enables researchers to describe, predict, and estimate the relationships between one or more independent variables and a dependent variable and draw plausible conclusions. Moreover, catterplots are one of the most common types of graphs that depicts the relationships between two variables. A nice scatterplot conveys the correlation, the spread, and outliers at the same time.

The case study demonstrates beginners in data analysis with cross-section data by estimating the Ordinary Least Square (OLS) regression, with panel data (longitudinal data) that contains observations about different cross sections across time by conducting panel regression, and with a nice two-way scatterplot.

In the multivariate regressions, the estimated coefficients for median household income are positive but statistically insignificant. When looking solely at the two-way scatterplot, median household income is positively associated with high school graduation rates.

**Coding language(s)**

Stata

Prompts to d*efine chatbot’s behaviors*

Act as an experienced economist proficient in Stata, you will write code with a number of lines as minimal as possible.

The input file(s) description will be [like this].

Reply “Input received” when receiving text with [] and explain.

Details of data analysis will be {like this}.

Reply “Analysis request received” when receiving text within {}, explain the text, followed by generating the code.

Reply “Yes” if understood.

### INPUT FILE

**Prompts**

[I have an Excel file: County Level Data.xls. I have variable names in the first row of my worksheet. The variable names are CountyID, year, HighSchoolGraduationRate, MedianHouseholdIncome, IncomeInequalityRatio, and SingleParentHouseholdRatio.]

**Code**

N/A

**Results**

N/A

**Assessments**

N/A

**REQUEST 1**

**Prompts**

{Run an OLS regression of HighSchoolGraduationRate on MedianHouseholdIncome, IncomeInequalityRatio, and SingleParentHouseholdRatio.}

**Code**

import excel "County Level Data.xls", sheet("Sheet1") firstrow clear

reg HighSchoolGraduationRate MedianHouseholdIncome IncomeInequalityRatio SingleParentHouseholdRatio

**Result**

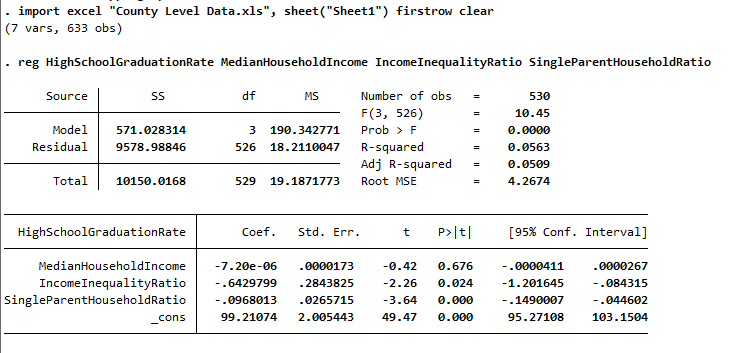

**Assessment**

The code runs successfully.

The Excel file is successfully imported into Stata in numerical format.

The code to conduct the OLS regression analysis is correct.

### REQUEST 2

**Prompts**

{How about run a panel regression with year fixed effects?}

**Code**

import excel "County Level Data.xls", sheet("Sheet1") firstrow clear

xtset CountyID year

xtreg HighSchoolGraduationRate MedianHouseholdIncome IncomeInequalityRatio SingleParentHouseholdRatio i.year, fe

**Result**

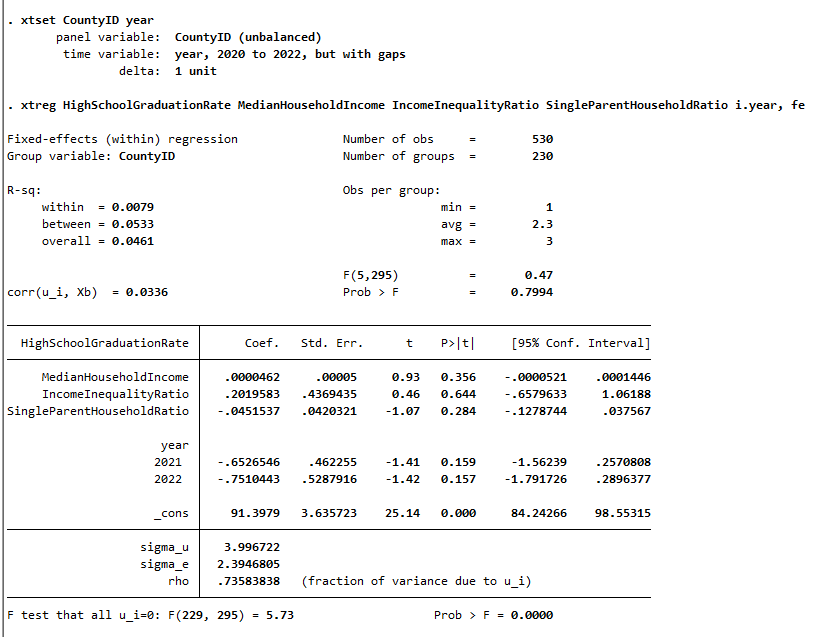

**Assessment**

The code run successfully. The panel regression analysis is completed.

### REQUEST 3 (ITERATION 0)

**Prompts**

{Show the relationship between HighSchoolGraduationRate and MedianHouseholdIncome in a figure}

**Code**

import excel "County Level Data.xls", sheet("Sheet1") firstrow clear

tw scatter HighSchoolGraduationRate MedianHouseholdIncome, xtitle("Median Household Income") ytitle("High School Graduation Rate") graphregion(margin(large)) name(scatterplot, replace)

**Result**

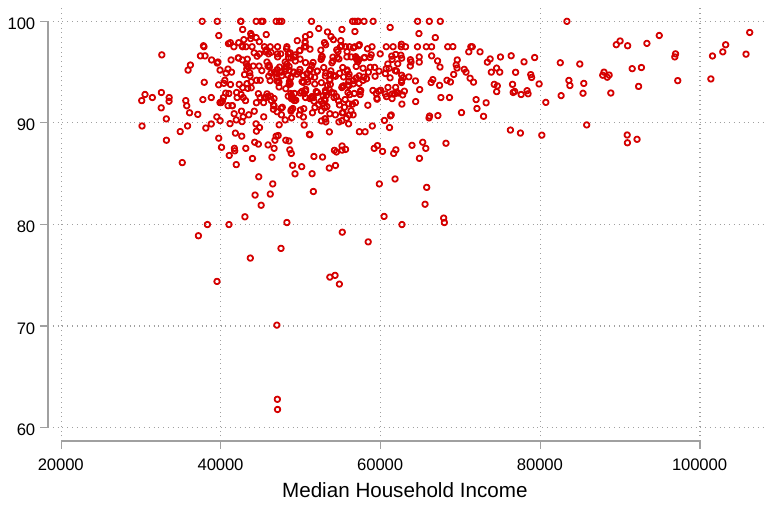

**Assessment**

The code run successfully.

The two-way scatterplots display the joint distribution of the HighSchoolGraduationRate and MedianHouseholdIncome variables.

### REQUEST 3 (ITERATION 1)

**Prompt**

{Add a title "High School Graduation Rate and Median House hold Income", a subtitle "County Level Evidence from Texas, 2020-2022", and a regression line and in the figure.}

**Code**

import excel "County Level Data.xls", sheet("Sheet1") firstrow clear

regress HighSchoolGraduationRate MedianHouseholdIncome

twoway (scatter HighSchoolGraduationRate MedianHouseholdIncome, xtitle("Median Household Income") ytitle("High School Graduation Rate") graphregion(margin(large)) name(scatterplot, replace)) (lfit HighSchoolGraduationRate MedianHouseholdIncome), title("High School Graduation Rate and Median Household Income") subtitle("County Level Evidence from Texas, 2020-2022")

**Result**

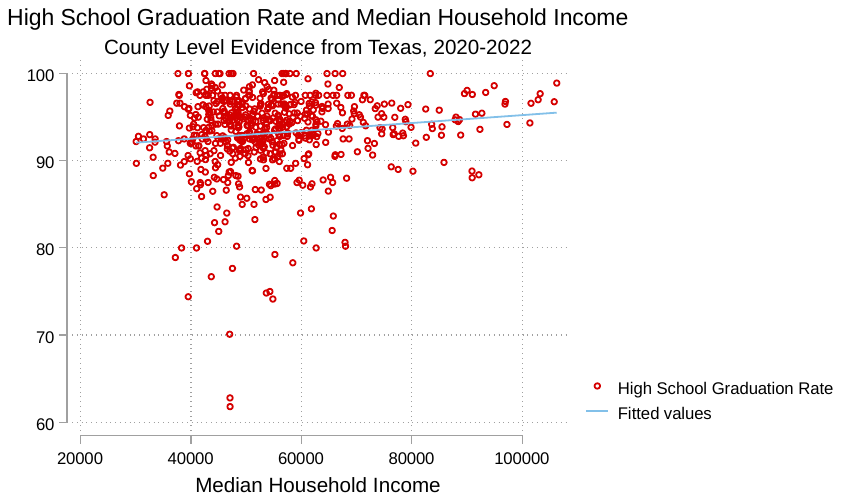

**Assessment**

The code for the OLS regression is unnecessary.

All other codes are good.

### WRAPPING UP

**Review the final code**

Review the comments added by chatGPT and identify key steps, as represented by functions and parameters, that are missing in the initial prompts.

**Prompts**

{I need to insert line-by-line comments to the below code which works well to address the needs for the data analysis task. Wait for my code.}

import excel "County Level Data.xls", sheet("Sheet1") firstrow clear

reg HighSchoolGraduationRate MedianHouseholdIncome IncomeInequalityRatio SingleParentHouseholdRatio

xtset CountyID year

xtreg HighSchoolGraduationRate MedianHouseholdIncome IncomeInequalityRatio SingleParentHouseholdRatio i.year, fe

twoway (scatter HighSchoolGraduationRate MedianHouseholdIncome, xtitle("Median Household Income") ytitle("High School Graduation Rate") graphregion(margin(large)) name(scatterplot, replace)) (lfit HighSchoolGraduationRate MedianHouseholdIncome), title("High School Graduation Rate and Median Household Income") subtitle("County Level Evidence from Texas, 2020-2022")

**Code with explanations**

// Import the data from the "County Level Data.xls" file

// with the variable names in the first row of the worksheet

import excel "County Level Data.xls", sheet("Sheet1") firstrow clear

// Run a simple linear regression of HighSchoolGraduationRate on MedianHouseholdIncome,

// IncomeInequalityRatio, and SingleParentHouseholdRatio

reg HighSchoolGraduationRate MedianHouseholdIncome IncomeInequalityRatio SingleParentHouseholdRatio

// Set the data as a panel with CountyID as the panel variable and year as the time variable

xtset CountyID year

// Run a fixed effects panel regression of HighSchoolGraduationRate on MedianHouseholdIncome,

// IncomeInequalityRatio, and SingleParentHouseholdRatio, with year fixed effects

xtreg HighSchoolGraduationRate MedianHouseholdIncome IncomeInequalityRatio SingleParentHouseholdRatio i.year, fe

// Create a scatter plot of HighSchoolGraduationRate against MedianHouseholdIncome

// with x and y axis titles and a large graph region, and save it as "scatterplot"

// Then, overlay a linear regression line on top of the scatter plot and add a title

// and subtitle to the graph

twoway (scatter HighSchoolGraduationRate MedianHouseholdIncome, xtitle("Median Household Income") ytitle("High School Graduation Rate") graphregion(margin(large)) name(scatterplot, replace)) (lfit HighSchoolGraduationRate MedianHouseholdIncome), title("High School Graduation Rate and Median Household Income") subtitle("County Level Evidence from Texas, 2020-2022")

### SUMMARY

**Aims**

The goal of this session is to input data in the Excel format in the statistical software and then run regression analysis and create an intutitive graph.

**Methods**

By providing appropriate prompts to ChatGPT, the case study demonstrates how beginners could obtain code from ChatGPT to run OLS regression for cross-section data and panel regression with time fixed effect for longitudinal data, along with data scatter plots.

**Results and Discussions**

Regression analysis is probably the oldest and most widely adopted quantitative data analysis technique in the social sciences. With the statistical method used to evaluate the relationship between the dependent variable and independent variables, beginners in data analysis can understand the significance of their data points and use analytical techniques to make better decisions.

The pattern of dots on a scatterplot, along with a fitted line for a simple [regression](https://statisticsbyjim.com/glossary/regression-analysis/) model, is a frequently used and effective way to illustrate the relationship or [correlation](https://statisticsbyjim.com/glossary/correlation/) between two continuous variables.

The codes obtained from ChatGPT are satisfactory. Besides providing the codes, ChatGPT also gives a detailed explanation. The process dramatically helps beginners learn how to interact with ChatGPT to resolve their demands, such as inputting data and conducting analysis, in a short time.

**Additional Comments**

Students need to know the subject matter to craft the prompts and ask pertinent questions. Without clear instructions on the object of the analysis, ChatGPT will have difficulty providing the correct code.
