## Supplementary material for "Empowering Beginners in Bioinformatics with ChatGPT": TableS5

**OPTICAL CASE STUDY: Time Series Analysis of Financial Data**

**Description***

The initial step in conducting a time series analysis typically involves computing and graphing the returns of a financial asset. For this particular case study, we will be utilizing actual trading price data for one-month maturity crude oil futures. Our goal is to determine the crude oil futures returns over time and pinpoint the date of any event that may have caused a significant price shock. This is a fundamental task in any financial data analysis.

We expect to obtain a visual representation of the daily return trend over a certain period. With this, we can move on to more advanced analyses such as generating summary statistics, plotting the data's histogram and comparing it to a normal distribution, identifying events linked to substantial negative returns, conducting Value at Risk analysis, forecasting conditional volatility, and performing portfolio analysis, among others.

**Coding language(s)**

Matlab

Prompts to d*efine chatbot’s behaviors*

Act as an experienced researcher in finance who is proficient in Matlab, you will write code with a number of lines as minimal as possible. The description of the input file(s) will be [like this]. Reply “Received input variables”. When a data analysis request is submitted, explain the text, followed by generating the code.

% Inputs:

[F:\Research2023\ChatGPT\Crude_Oil_Data.csv]

% Reply:

Input received

**ITERATION 0**

**Prompts**

Read crude oil futures price data; calculate and plot log returns of crude oil futures prices; circle the lowest return and mark it with the date.

**Code**

% Read the crude oil futures price data

data = readtable('F:\Research2023\ChatGPT\Crude_Oil_Data.csv');

% Calculate log returns of crude oil futures prices

log_returns = diff(log(data.Price));

% Find the lowest return and its date

[lowest_return, index] = min(log_returns);

date_of_lowest_return = data.Date(index + 1);

% Plot the log returns of crude oil futures prices

plot(log_returns);

hold on;

% Circle the lowest return and mark it with the date

plot(index, lowest_return, 'ro', 'MarkerSize', 10, 'LineWidth', 2);

text(index, lowest_return, date_of_lowest_return, 'HorizontalAlignment', 'center', 'VerticalAlignment', 'top');

hold off;

% Add title and axis labels

title('Log Returns of Crude Oil Futures Prices');

xlabel('Time');

ylabel('Log Returns');

**Result**

Unrecognized table variable name 'Price'.

**Assessment***

The code failed to recognize a variable name and cannot load the variable.

**ITERATION 1**

**Prompts**

Unrecognized table variable name 'Price'.

**Response**

I'm sorry for the error. It looks like the variable name for the crude oil futures price column in the input file is different. Please provide me with the column name for crude oil futures price data in your input file, and I'll adjust the code accordingly.

**Assessment***

It seems that ChatGPT has problems reading some variable names correctly. It needs human intervention.

The variable name in the original data was “Crude Oil Futures Price” in the input csv file. When it was loaded into Matlab, it was “CrudeOilFuturesPrice,”

In a previous trial, ChatGPT once read the variable name as “CrudeOilFutures.”

When fed with the prompt: “Unrecognized table variable name ' CrudeOilFutures'”, ChatGPT responded and suggested replacing this line:

crude_oil_prices = crude_oil_data.CrudeOilFutures;

with this line:

crude_oil_prices = crude_oil_data.Crude_Oil_Futures;

This still didn’t solve the problem and couldn’t load the proper variable.

We have to manually fix it using two alternatives. One way is to tell ChatGPT that the variable name should be “CrudeOilFuturesPrice”; the other way is to change the variable name in the original csv file to “Price” (use a variable name without spaces).

For the rest of the discussion, we used the second one, changing the variable name in the original input file to “Price.”

**ITERATION 2**

**Prompts**

Read crude oil futures price data; calculate and plot log returns of crude oil futures prices; circle the lowest return and mark it with the date.

**Code**

% Read the crude oil futures price data

data = readtable('F:\Research2023\ChatGPT\Crude_Oil_Data.csv');

% Calculate log returns of crude oil futures prices

log_returns = diff(log(data.Price));

% Find the lowest return and its date

[lowest_return, index] = min(log_returns);

date_of_lowest_return = data.Date(index + 1);

% Plot the log returns of crude oil futures prices

plot(log_returns);

hold on;

% Circle the lowest return and mark it with the date

plot(index, lowest_return, 'ro', 'MarkerSize', 10, 'LineWidth', 2);

text(index, lowest_return, date_of_lowest_return, 'HorizontalAlignment', 'center', 'VerticalAlignment', 'top');

hold off;

% Add title and axis labels

title('Log Returns of Crude Oil Futures Prices');

xlabel('Time');

ylabel('Log Returns');

**Result**

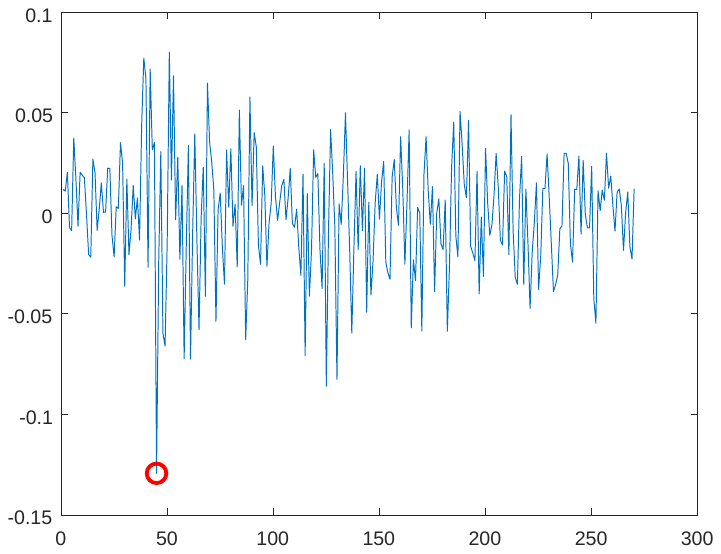

**Assessment***

The data was loaded successfully, and the time series of calculated log returns was plotted. However, the label for x-axis does not show the date information. There are no label for axis or figure title. The lowest return was circled but didn’t show its date.

It also gives an error message when adding the text next to the lowest returns (see below).

Error using text

Invalid parameter/value pair arguments

Error in VaR_test3 (line 17)

text(index, lowest_return, date_of_lowest_return, 'HorizontalAlignment', 'center', 'VerticalAlignment', 'top');

We need to give ChatGPT more specific commands.

**ITERATION 3**

**Prompts**

Plot log returns of crude oil futures prices as a function of date; show label for the axis and figure title; circle the lowest return and mark it with its date.

**Code**

% Read the crude oil futures price data

data = readtable('F:\Research2023\ChatGPT\Crude_Oil_Data.csv');

% Calculate log returns of crude oil futures prices

log_returns = diff(log(data.Price));

% Find the lowest return and its date

[lowest_return, index] = min(log_returns);

date_of_lowest_return = data.Date(index + 1);

% Plot the log returns of crude oil futures prices as a function of date

plot(data.Date(2:end), log_returns);

hold on;

% Circle the lowest return and mark it with the date

plot(date_of_lowest_return, lowest_return, 'ro', 'MarkerSize', 10, 'LineWidth', 2);

text(date_of_lowest_return, lowest_return, num2str(lowest_return), 'HorizontalAlignment', 'center', 'VerticalAlignment', 'top');

hold off;

% Add title and axis labels

title('Log Returns of Crude Oil Futures Prices');

xlabel('Date');

ylabel('Log Returns');

% Add labels for the y-axis rises

ax = gca;

ax.YAxis.Exponent = 0;

ax.YRuler.Exponent = 0;

ytickformat('%.2f');

**Result**

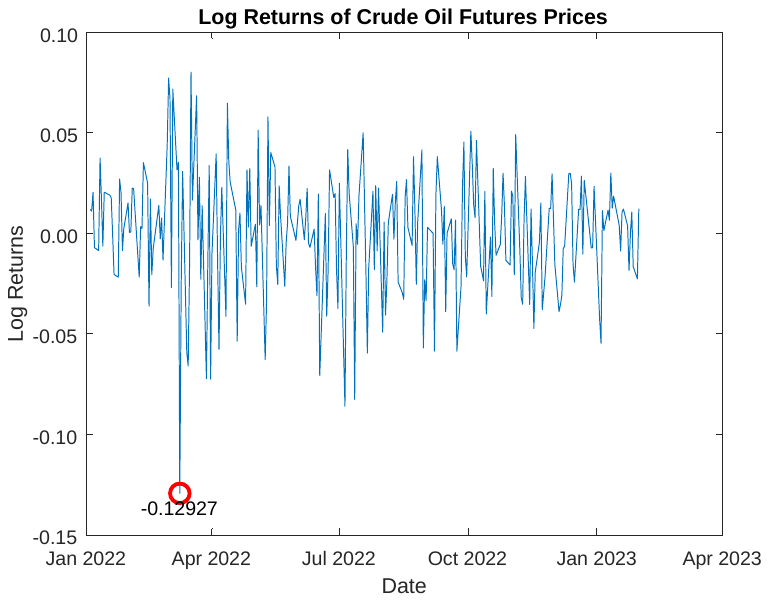

**Assessment***

The returns are plotted as directed, with date, y label, and figure title. Now the only problem is that the circled lowest return was marked by its log return value, not by its date. We want to identify the date so as to link it with the event on that date to interpret the lowest return. Therefore, we need to emphasize that we want the lowest return marked with the date, not the return.

**ITERATION 4**

**Prompts**

For the lowest return, please add a text next to it. The text should be the date associated with this return, not the log return value.

**Code**

% Read the crude oil futures price data

data = readtable('F:\Research2023\ChatGPT\Crude_Oil_Data.csv');

% Calculate log returns of crude oil futures prices

log_returns = diff(log(data.Price));

% Find the lowest return and its date

[lowest_return, index] = min(log_returns);

date_of_lowest_return = data.Date(index + 1);

% Plot the log returns of crude oil futures prices as a function of date

plot(data.Date(2:end), log_returns);

hold on;

% Find the crude oil futures price at the date of the lowest return

price_of_lowest_return = data.Price(index + 1);

% Circle the lowest return and mark it with the date

plot(date_of_lowest_return, log_returns(index), 'ro', 'MarkerSize', 10, 'LineWidth', 2);

text(date_of_lowest_return, log_returns(index), datestr(date_of_lowest_return), 'HorizontalAlignment', 'center', 'VerticalAlignment', 'bottom');

hold off;

% Add title and axis labels

title('Log Returns of Crude Oil Futures Prices');

xlabel('Date');

ylabel('Log Returns');

% Add labels for the y-axis rises

ax = gca;

ax.YAxis.Exponent = 0;

ax.YRuler.Exponent = 0;

ytickformat('%.2f');

**Result**

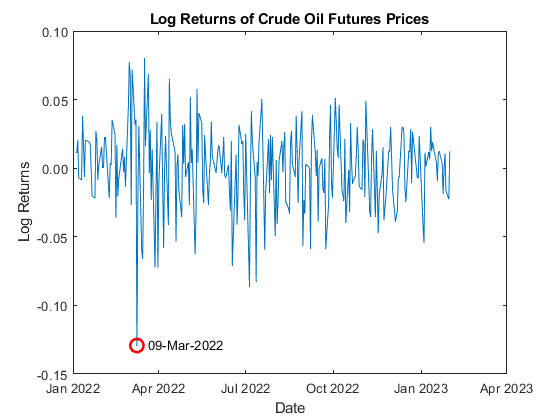

**Assessment***

The above plot shows the expected results after a few attempts.

However, to get the ideal plot, this prompt has to be very detailed and clear. I have tried a few other ways that didn’t work. For example, “Please mark the lowest return with its corresponding date,” or “Please mark the lowest return in the plot with its corresponding date only, not with its log return value.”

**WRAPPING UP**

**Review the final codes**

**Prompts**: “Include line-by-line explanation in the immediately previous code”

Here's the updated code with line-by-line explanations:

% Read the crude oil futures price data

data = readtable('F:\Research2023\ChatGPT\Crude_Oil_Data.csv');

% Reads the data from the specified file and stores it in a table named 'data'

% Calculate log returns of crude oil futures prices

log_returns = diff(log(data.Price));

% Calculates the log returns of crude oil futures prices using the 'diff' and 'log' functions. The resulting values are stored in a column vector named 'log_returns'.

% Find the lowest return and its date

[lowest_return, index] = min(log_returns);

% Determines the minimum value in the 'log_returns' vector and its index, which correspond to the lowest return and the date of the day after the day with the lowest return, respectively. These values are stored in 'lowest_return' and 'index'.

date_of_lowest_return = data.Date(index + 1);

% Finds the date of the day after the day with the lowest return using the 'index' variable and the 'Date' column of the 'data' table. The resulting date is stored in 'date_of_lowest_return'.

% Plot the log returns of crude oil futures prices as a function of date

plot(data.Date(2:end), log_returns);

% Plots the log returns of crude oil futures prices using the 'plot' function. The 'Date' column of the 'data' table (excluding the first value) is used as the x-axis and the 'log_returns' vector is used as the y-axis.

hold on;

% Find the crude oil futures price at the date of the lowest return

price_of_lowest_return = data.Price(index + 1);

% Finds the crude oil futures price at the date of the day after the day with the lowest return using the 'index' variable and the 'Price' column of the 'data' table. The resulting price is stored in 'price_of_lowest_return'.

% Circle the lowest return and mark it with the date

plot(date_of_lowest_return, log_returns(index), 'ro', 'MarkerSize', 10, 'LineWidth', 2);

% Plots a red circle at the position of the day after the day with the lowest return using the 'plot' function. The circle is centered at the date stored in 'date_of_lowest_return' and the y-coordinate stored in 'log_returns(index)'. The circle has a marker size of 10 and a line width of 2.

text(date_of_lowest_return, log_returns(index), datestr(date_of_lowest_return), 'HorizontalAlignment', 'center', 'VerticalAlignment', 'bottom');

% Adds a text label next to the circle using the 'text' function. The text is the date stored in 'date_of_lowest_return' and is centered at the same position as the circle. The horizontal alignment is centered and the vertical alignment is at the bottom.

hold off;

% Add title and axis labels

title('Log Returns of Crude Oil Futures Prices');

% Adds a title to the plot using the 'title' function.

xlabel('Date');

% Adds a label to the x-axis using the 'xlabel' function.

ylabel('Log Returns');

% Adds a label to the y-axis using the 'ylabel' function.

% Add labels for the y-axis rises

ax = gca;

% Gets the current axis using the 'gca' function and stores it in 'ax'.

ax.YAxis.Exponent = 0;

% Sets the y-axis exponent to zero, which removes scientific notation for the y-axis tick labels.

ax.YRuler.Exponent = 0;

% Sets the y-axis ruler exponent to zero, which

**Summary***

**Aims**

The objective of this study is to prepare the crude oil futures data for further financial analysis. To achieve this, we calculate and graph the log returns of crude oil futures prices, and determine the event of a price shock.

This study marks the initial stage in financial time series analysis that numerous empirical analyses would expand upon.

**Methods**

The methods used in this study are fundamental in financial analysis. To conduct time-series analysis, we computed the log returns of crude oil futures prices using the formula $R_{t+1}=ln(\frac{P_{t+1}}{P_{t}})$, where $R_{t}$ is the log return and $P_{t}$ is the crude oil futures price. This method is a widely accepted practice in financial analysis and is useful for identifying trends and patterns in market behavior, which can help inform investment decisions.

**Results and Discussions**

In general, the results are satisfactory, but there were some issues that ChatGPT encountered, such as reading the variable name and marking the date of a price shock event. However, these difficulties presented valuable opportunities for students to learn how to interact with ChatGPT effectively, such as selecting appropriate variable names and refining prompts with more specific commands to achieve the desired outcome. These practices can help enhance their coding skills and analytical abilities, which can be beneficial in various fields, including finance and data analysis.

**Additional Comments***

We sent ChatPGT a revised final prompt in a new chat window: “Read crude oil data; calculate returns of crude oil futures prices (variable name “Price” in the data); Plot log returns of crude oil futures prices as a function of date; show label for the axis and figure title; circle the lowest return and add a text next to it. The text should be the date associated with this return, not the log return value.” The code generated by ChatGPT successfully produced the desired results.

From this study, we learned the importance of providing simple and specific instructions and evaluating results step by step. Additionally, we noticed that ChatGPT sometimes provides irrelevant information from past interactions, but generally performs well with clear instructions.

*The writing has been polished by ChatGPT after an initial human draft.
